# Rapid and highly sensitive detection of pyocyanin biomarker in different *Pseudomonas aeruginosa* infections using gold nanoparticles modified sensor

**DOI:** 10.1101/616797

**Authors:** Amal A. Elkhawaga, Marwa M. Khalifa, Omnia H.B. El-badawy, Mona A. Hassan, Waleed A. El-Said

## Abstract

Successful antibiotic treatment of infections relies on accurate and rapid identification of the infectious agents. *Pseudomonas aeruginosa* is implicated in a wide range of human infections that almost complicated and become life threating especially in immunocompromised and critically ill patients. Conventional microbiological methods take more than 3 days to obtain accurate results. Pyocyanin is a distinctive electroactive biomarker for *Pseudomonas aeruginosa*. Here, we have developed a rapid diagnostic (polyaniline) PANI gold nanoparticles (Au NPs) modified indium tin oxide (ITO) electrode that showed 100% sensitivity for pyocyanin in culture of *Pseudomonas aeruginosa* clinical isolates and high selectivity for pyocyanin at low concentration when measured in the presence of other substances like ascorbic acid, uric acid, and glucose as interferences. The constructed electrode was characterized using scanning electron microscopy and cyclic voltammetry. The determined linear range for pyocyanin detection was from 238 µM to 1.9 µM with a detection limit of 500 nM. Compared to the screen-printed electrode used before, the constructed electrode showed a 4-fold enhanced performance.

## 1. Introduction

*Pseudomonas aeruginosa* (*P. aeruginosa*) is a prevalent and opportunistic pathogen that is considered one of the most annoying bacteria causing deadly infections in critically ill patients [**1-4**]. It commonly produces infections in patients with surgical wounds, burn wound or cystic fibrosis. Infections caused by *P. aeruginosa* have high morbidity and mortality rates, particularly among immunocompromised patients such as cancer patients and premature infants [**5-7**]. *P. aeruginosa* may acquire multidrug resistance, making its eradication with antibiotics challenging [**8**]. The increasing resistance of bacteria is partially due to the late diagnosis and the misuse of antibiotics [**9**]. Hence, the early and fast detection of this serious pathogen is essential for a more targeted antibiotic prescription that will hasten recovery and reduce the emergence of antibiotic resistance [**10, 11**]. Typically, *P. aeruginosa* infections are identified in clinics using selective plate culturing techniques, which take about 24 h or more to provide results. Polymerase chain reaction (PCR) identification is available in a growing number of clinics. However, PCR identification requires extensive sample preparations, uses expensive reagents, and takes several hours to complete [**12**]. Therefore, developing a sensitive, specific and rapid identification methods for pathogen detection in cost and time competent manners have found broad attention in the last few years [**13**]. Pyocyanin is one of the virulence factors exclusively secreted by *P. aeruginosa*. It is a unique, quorum-sensing molecule that is linked to biofilm formation, induces inflammation and causes apoptosis of neutrophils [**14-16**].

Until lately, pyocyanin was measured by chromatography or spectrophotometric techniques that are time-consuming and needs purification from bacterial cultures [**15**]. Electrochemical sensors are endorsed robust tools for the detection of environmental and disease-related biomarkers [**13**]. They are easy to use with low detection limits, high sensitivity, good stability together with cost and time effectiveness [**17, 18**]. The redox-active nature of pyocyanin molecules permits its rapid detection by electrochemical sensors in two minutes [**19**]. It is still a challenge to find new and more sensitive methods to provide rapid and accurate information about *P. aeruginosa* that can aid treatment decisions as early as possible when bacteria are still responsive to antibiotics.

Gold (Au) nanostructures are currently used to modify electrodes of biosensors because of its excellent optical and electrical properties and affinity to bind with biomolecules [**20-25**]. Moreover, the use of Au modified indium tin oxide (ITO) electrode has led to a fast and ultrasensitive detection of some multidrug-resistant bacteria such as *E. coli* and *S. aureus* [**26**].

Conducting polymers [**27**] have much interest in the recent researchers because of their good conductivity, stability, ease of preparation. In general, the electronic and electrochemical properties of [**28-31**] conducting polymers made them have many applications in photovoltaic cells, organic light emitting diode, and biosensors. Polyaniline has received much attention in the research work. This is mainly due to the fact that PANI and its derivatives or composites with other materials are easy to synthesize chemically or electrochemically [**32**].

Therefore, conducting polymers/metal or metal oxides hybrid materials possess the unique combination of the conducting polymers properties (biocompatibility, direct electrochemical synthesis) and the characteristics of nanomaterials e.g. large surface area, size, flexibility for the immobilization of biomolecules and quantum effect [**33**]. In our previous work, we have reported on the fabrication of poly(4–aminothiophenol) nanostructures decorated gold nanodots patterned ITO electrode, which demonstrated a highly electrochemical sensitivity and selectivity towards a mixture of two DNA bases (adenine and guanine) [**24**].

Hybrids organic and inorganic nanocomposites not only possess the sum of their individual components, but the role of the inner interfaces could be predominant; thus hybrids organic/inorganic nanocomposites have used in a wide range of applications including biosensors [**22, 34-38**] and sensors [**39**].

The present work aims to assess the efficacy of using PANI/Au nanostructures modified ITO sensor for early detection and quantification of pyocyanin in *P. aeruginosa* cultures of clinical isolates based on cyclic voltammetry (CV) technique. The developed sensor showed high sensitivity towards pyocyanin over a wide range of concentrations from 238 µM to 1.9 µM with a limit of detection (LOD) of about 500 nM. Also, this sensor showed high selectivity towards detection of pyocyanin in the presence of several interferences such as urea, glucose and vitamin c. Furthermore, the applicability of this sensor has been confirmed by direct detection of pyocyanin in *P. aeruginosa* culture of clinical isolates obtained from cases of *P. aeruginosa* infections.

## 2. Methods

### 2.1. Materials

Pyocyanin (P0046-5MG), gold (III) chloride hydrate and ITO coated glass slide square were obtained from Sigma Aldrich. Deionized water (DIW) with a resistivity of 18.2 MΩ.cm that was purified with a Purite purification system (UK) was used for all preparations. The buffer system used in this work was phosphate buffer saline (PBS) (0.01 mol/L) at pH 7.4 that was prepared by dissolving PBS powder in 1 L of DIW. Luria-Bertani (LB) broth (Oxoid, UK) was utilized in this study.

### 2.2. Equipment

All electrochemical measurements were performed using the Autolab potentiostat instrument (Netherlands) connected to a three-electrode cell; Metrohm Model 663VA stand was controlled by Nova software at room temperature. The three-electrode system consists of a platinum wire as a counter electrode, Ag/AgCl as the reference electrode, and PANI/Au modified ITO electrode as a working electrode.

### 2.3. Clinical isolates of Pseudomonas aeruginosa

*P. aeruginosa* culture was made from clinical isolates obtained from the department of Medical Microbiology and Imunology that was isolated from clinical cases of *P. aeruginosa* infections admitted to Assiut University hospital as pneumonia, corneal ulcers, urinary tract infections and wound infections. These clinical isolates of *P. aeruginosa* were confirmed by VITEK 2 automated microbiology system. The study protocol was approved by the local ethical Committee of the faculty of Medicine Assiut University and an informed written consent was taken from all the participants in the study.

### 2.4. Methods

#### 2.4.1. Preparation of gold nanostructures ITO electrode

The ITO-coated glass substrates with a geometrical size of 25 mm X 12.5 mm X 1.1 mm was cleaned *via* sonication for 15 min each in 1% Triton X-100, DIW and then in ethanol. The substrates were immersed in a basic piranha solution (H_2_O_2_: NH_3_: H_2_O ratio, 1:1:5) for 30 min at 80 °C. Finally, the substrates were cleaned again with DIW and ethanol and dried under nitrogen gas. The Au NPs modified ITO electrodes were prepared according to our previously reported method [**21**] in which an aqueous solution of 1 mM HAuCl_4_ was added into the electrochemical cell and we have issued the deposition process of Au NPs on the ITO substrates by using CV technique within potential window from 1.5 V to −1 V for 5 cycles at scan rate of 50 mV/sec against Ag/AgCl as a reference electrode. The surface morphology of the modified electrodes was analyzed by SEM (JOEL-JSM-5400LV).

#### 2.4.2. Preparation of gold modified ITO electrode

Polyaniline hydrochloride (PANI) salt was prepared according to the previously published work [**40**]. Typically, 0.5 g of aniline hydrochloride was dissolved in 20 mL of DIW and stirred in ice bath for 1h (first solution). In another conical flask, a solution of 0.5 g of ammonium persulphate in 20 mL DIW was stirred in ice bath for 1h and then added to the first solution and keep stirring for further 4h. The dark green precipitates were filtrated and dried in an oven at 80 °C [**41**]. To fabricate a layer of PANI on the surface of Au NPs/ITO electrode, Au NPs/ITO electrode was immersed in a solution of PANI in NMP (0.001 gm/mL) for 8 hrs and then rinsed with DIW to remove the PANI from the non-conductive side and dried under N_2_ gas [**40, 42**].

#### 2.4.3. Electrochemical Measurements of pyocyanin

The electrochemical measurements were carried out by immersing the working electrode together with the reference and counter electrodes in the presence of different concentrations of pyocyanin ranging from 238 µM to 1.9 µM; the solutions were prepared in PBS (10 mmol/L, pH 7.4).

#### 2.4.4. Selectivity of the Developed pyocyanin Sensor

The selectivity of the prepared electrodes towards the pyocyanin was studied by using a mixture of pyocyanin solution with glucose, vitamin c, and urea as interferences, which are commonly present in clinical samples.

#### 2.4.5. Electrochemical detection of pyocyanin in Pseudomonas aeruginosa

Under complete sterile conditions a colony (or more) of *P. aeruginosa* were added in 10 ml of LB broth in 14 ml tube and placed on a shaker (200 rpm) at 37 °C overnight. 1ml of this suspension was removed and added to 9 ml fresh LB broth in 14 ml tube and placed again in a shaker at 37 °C for 24 hrs. The OD at 600 nm (OD600) was measured to quantify the density of the bacteria in each culture sample. During the 24 hours, 3 samples were collected at different time-points (after 2, 10 and 24 hours). The pyocyanin concentration was measured using the PANI/Au NPs modified ITO electrode and the OD600 was detected to confirm the increasing bacterial number.

#### 2.4.6. Electrochemical testing of bacterial cultures

Each strain of *P. aeruginosa, Staphylococcus aureus, Staphylococcus epidermidis, Klebsiella pneumoniae and Streptococcus pneumoniae* was cultivated in LB broth at 37 °C overnight. The cyclic voltammetry of each strain culture was measured using the PANI/Au NPs modified ITO electrode.

#### 2.4.7. Electrochemical detection of P. aeruginosa isolated from clinical samples

Sterile swabs were used to collect different *P. aeruginosa* clinical isolates for electrochemical testing. Swab samples were inoculated into LB broth and incubated at 37 °C overnight. CV of each sample was measured using the PANI/Au NPs modified ITO electrode.

## 3. Results and discussion

### 3.1 Preparation and Structural Features of the Au NPs/ITO and PANI/Au NPs/ITO Electrodes

One of the most critical advantages of using Au NPs for the preparation of biosensors is their ability to give a stable immobilization of biomolecules, thereby retaining their bioactivity. Moreover, Au NPs can allow electrons to be transported without using an electron transfer mediator by encouraging the electron transfer between redox proteins and electrode materials. Electron transfer is facilitated because Au NPs possess attractive characteristics, such as high surface-to-volume ratio, high surface energy, the ability to decrease protein-metal particle distance and to function as electron-conducting pathways [**43**].

In this work, Au NPs modified ITO electrode was prepared based on electrochemical deposition of Au onto the ITO surface by using CV technique. Figure 1 showed the cyclic voltammograms corresponding to the electrochemical deposition process within a potential window from 1.5 V to −1.0 V to allow a complete reduction of Au^3+^ ions into Au^0^ under this potential window range after 5 cycles [**20**]. **Figure 1** demonstrated a reduction peak at −0.5, −0.13 and 0.1 V, anodic peak at −0.1 and 0.84 V during the first cycle of Au nanoparticles deposition, which is shifted to −0.35 and 0.41 V and anodic peaks at 0.061 and 0.84 V during the deposition process in the remaining 4 cycles. The shifting in the reduction peaks is related to the reduction process of Au^3+^ to form metallic Au nanostructures on the ITO electrode surface [**20**].

**Figure 1.**
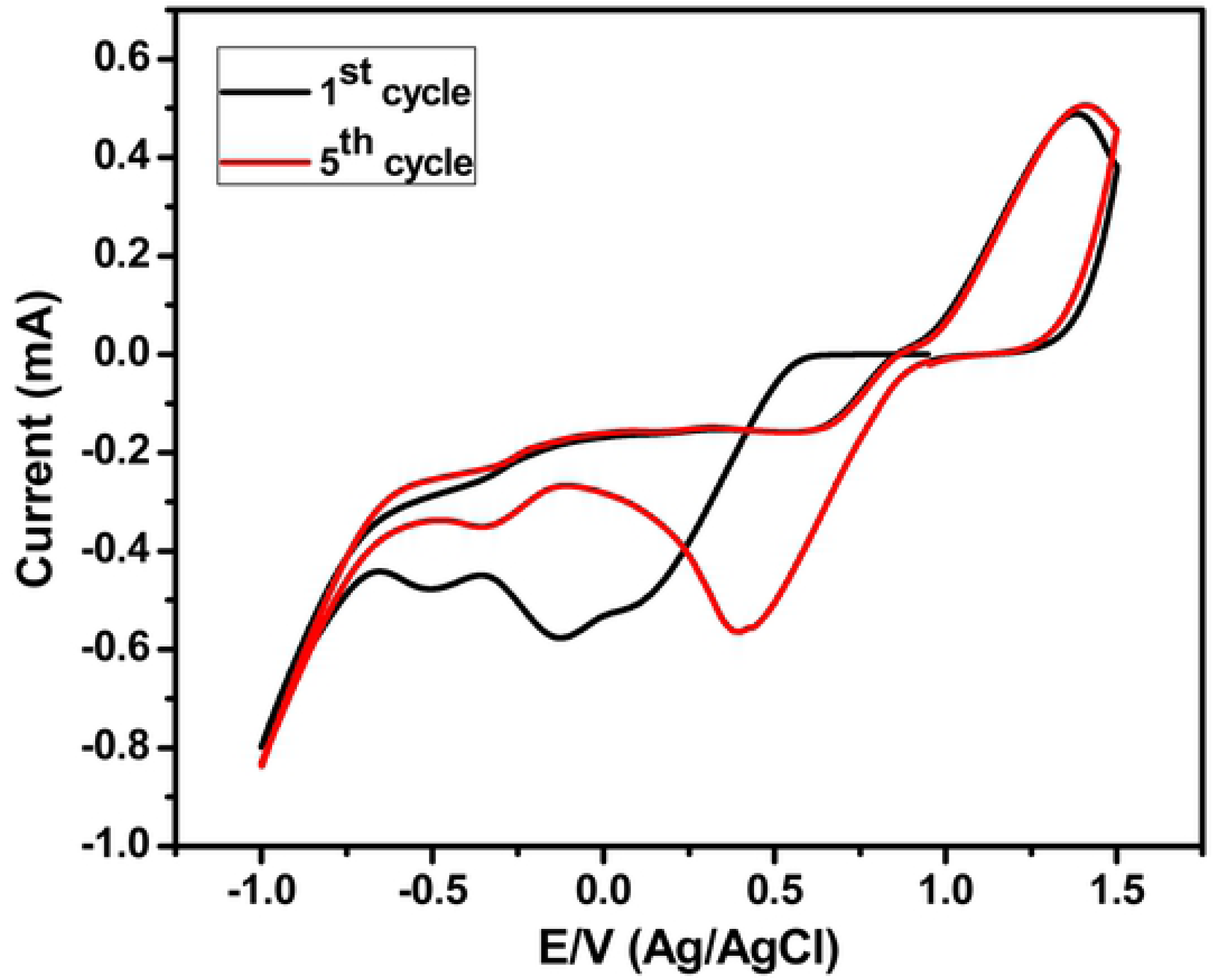
Electrochemical deposition of Au NPs onto ITO substrate based on CV technique within potential window from 1.5 V to −0.1 V. Scan rate is 50 mV/sec.

The nanostructured surface morphology of the Au NPs modified ITO electrode could also have a significant effect on the sensitivity of electrochemical detection, which was further explored by SEM characterization. As shown in **Figure 2a**, the SEM image of the Au nanostructured modified ITO electrode illustrated a nice coverage of Au NPs on the surface of the ITO electrode with the formation of polydispersed Au NPs. The particle size was analyzed by using SPIP (version 6.7.7) program (**Figure 2b**), which demonstrated that the mean particle size was found to be about 42.5 nm in diameter with a standard deviation of about 101 nm.

**Figure 2.**
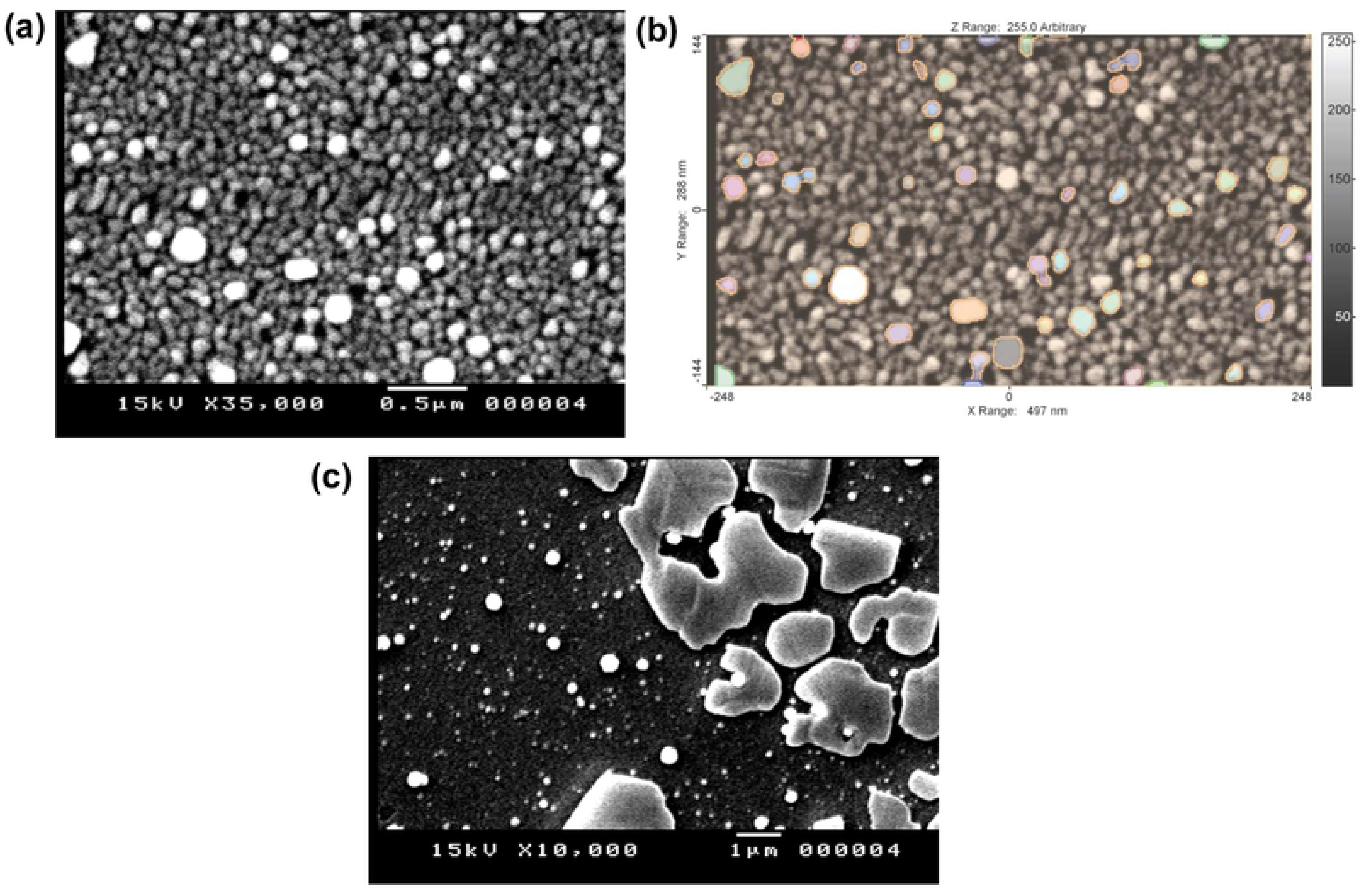
(a) SEM image of Au NPs modifed ITO electrode prepared after 5 cycles, (b) SPIP analysis of the SEM image of the Au NPs modifed ITO electrode prepared after 5 cycles and (c) SEM image of PANI/ Au NPs/ITO electrode.

### 3.2. The Electrochemical Behavior of pyocyanin by using Cyclic Voltammetry

**Figure 3a** showed the CV behavior of 50 µM of pyocyanin in PBS at the bare ITO electrode, which demonstrated a very weak redox peaks. So, the CV response of higher concentration was investigated (**Figure 3b**), which showed an anodic peak at −0.203 V and a cathodic peak at −0.305 V. Thus, the bare ITO electrode is unsuitable for detection low concentrations of pyocyanin. In order to develop an electrode that could sense the pyocyanin; we have modified the ITO electrode with Au NPs and used it to study the electrochemical behavior of pyocyanin based on CV technique. **Figure 3c** showed the cyclic voltammograms of three different concentrations of pyocyanin at Au NPs modified ITO electrode, which illustrated a quasirevisable response with an oxidation peak at −0.21 V and a reduction peak at about −0.3 V. These results indicate the capability of the Au NPs modified ITO electrode to detect the pyocyanin; this capability is attributed to the signal amplification of Au NPs that enhanced the electron transfer characterization [**21**]. However, the Au NPs modified ITO electrode didn’t show any response to pyocyanin solution with concentrations lower than 36 µM. To enhance the sensitivity of the developed electrode, we have modified the Au NPs/ITO electrode with a layer of PANI and used to study it to detect the pyocyanin marker. **Figure 2c** showed the SEM image of the PANI/Au NPs/ITO, which showed the formation of a thin layer of PANI with large diameter. The cyclic voltammogram of 50 µM pyocyanin at PANI/Au NPs/ITO electrode was represented in **Figure 3d**, which showed an increase in the oxidation peak at −0.23 V and reduction peak at −0.3 V. Furthermore, it is intersted to note that the redox current peak is higher than that in either case of using ITO or Au NPs/ITO electrodes, I addtion the revisabilty of the redox behavior was increased with electrode modification. So that PANI/Au NPs/ ITO electrode is more sensitive to pyocyanin than either bare ITO electrode or Au NPs/ITO electrode.

**Figure 3.**
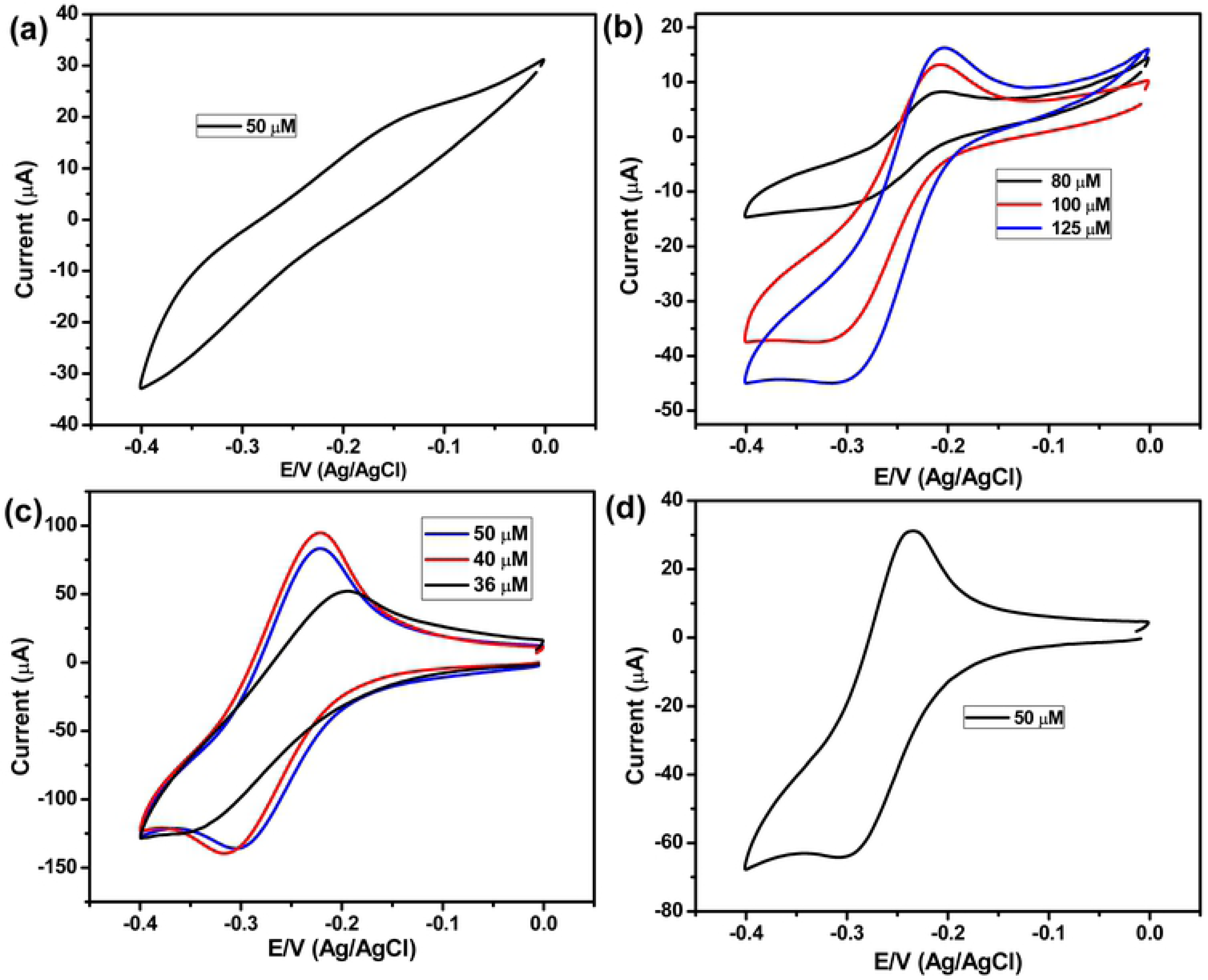
Cyclic voltammetry behavior of (a) 50 µM pyocyanin in PBS buffer at bare ITO, (b) three different concentrations of pyocyanin in PBS buffer at bare ITO, (c) three different concentrations of pyocyanin in PBS buffer at Au NPs modified ITO and (d) 50 µM pyocyanin in PBS buffer at PANI/Au NPs modified ITO. The scan rate was 50 mV/sec.

**Figure 4a** showed the effect of different scan rate within a range from 0.01 V/s to 0.12 V/s on the maximum peak current of pyocyanin, which illustrated an increase in the redox peaks with the increase of the scan rate. **Figure 4b** showed the relationship between the value of the scan rate versus the maximum peak current of pyocyanin at Au NPs/ITO electrode, which demonstrated a linear relationship over a wide range of scan rate from 0.01 V/s to 0.12 V/s.

**Figure 4.**
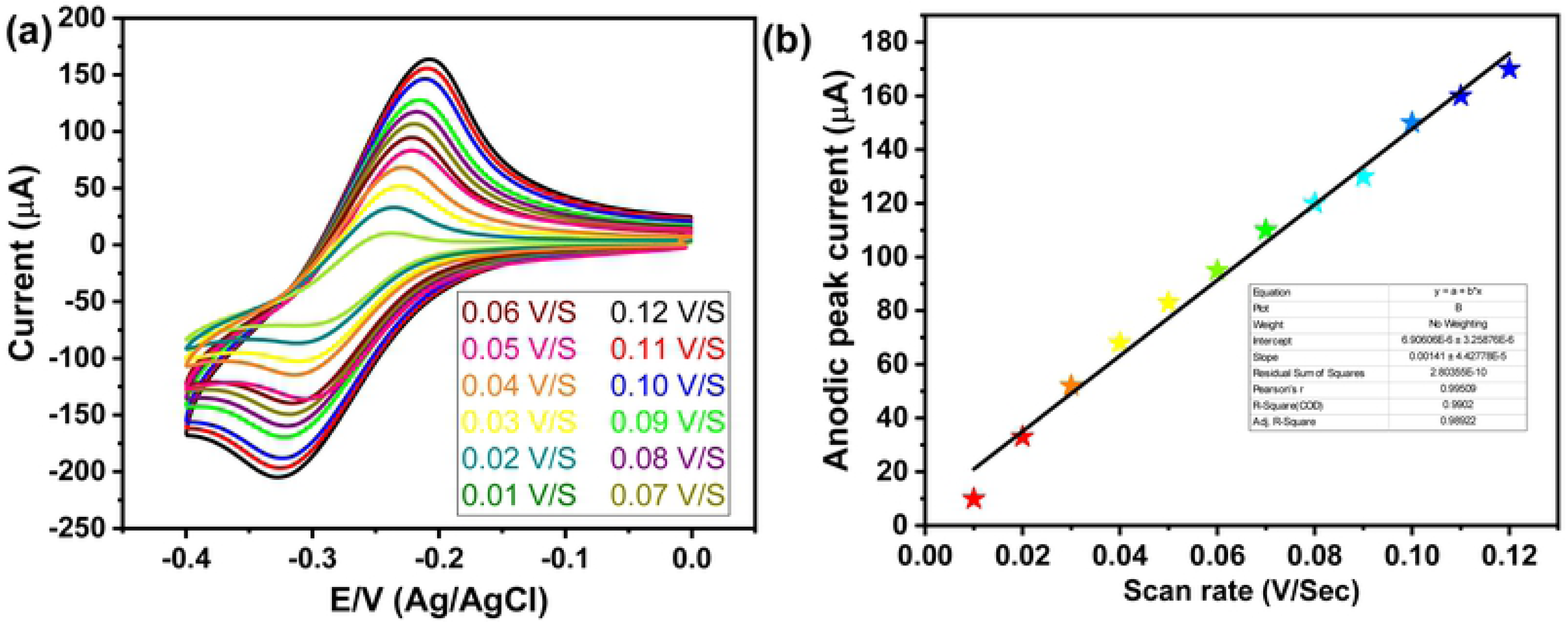
a) CV of pyocyanin 50 µM at different scan rate from 0.01 V/s to 0.12 V/s and b) scan rate versus the oxidation peak current of pyocyanin.

### 3.3. The sensitivity of the developed sensor towards pyocyanin marker

To evaluate the sensitivity of the PANI/Au NPs modified ITO electrode towards pyocyanin, a wide range of pyocyanin concentrations from 238 µM to 1.9 µM in PBS was used and their CVs response at PANI/Au NPs modified ITO electrode was investigated. **Figure 5a** showed the CVs bahvior of different concentrations of pyocyanin at PANI/Au NPs modified ITO electrode, which demonstrated an increase in the redox current peak with increasing the pyocyanin concentration. **Figure 5b** represented the relationship between the oxidation current peak and the pyocyanin concentration at PANI/Au NPs modified ITO electrode, which illustrated a linear response between the anodic current peaks and the concentration of pyocyanin. The LOD of the PANI/Au NPs modified ITO electrode was calculated according to the equation (LOD =3.3*(STEYX/Slope of calibration curve)), and it was found to be 500 nM. This result is better than the results obtained by Alatraktchi et al., who detected pyocyanin by a disposable screen-printed electrode [**44**]. The results in **Figure 5** showed the high sensitivity for detection of pyocyanin using the Au modified ITO, which attributed to the use of PANI/Au NPs modified ITO as a working electrode that achieved the precise, rapid and sensitive measurement of pyocyanin with low cost. **Table 1** showed the LOD of our modified electrode in comparison with the LOD of the previously reported of some other electrodes for the electrochemical determination of pyocyanin that revealed that our electrode possessed lower LOD in comparison with most of the previously reported electrodes [**44-50**].

**Table 1.** Comparison between the sensitivity of our sensor with the previous work

| **Sensor** | **Electrochemical technique** | **LOD** | **References** |
| --- | --- | --- | --- |
| Screen-printed electrode (gold working electrode) | CV | 2 $\mu\text{M}$ | (44) |
| Screen-printed electrode (gold working electrode) | Amperometry | 0.125 $\mu\text{M}$ | (45) |
| Screen printed sensing glove (carbon ink) | SWV | 0.003 $\mu\text{M}$ | (46) |
| Three electrode configurations consisting of a Carbon Fibre tow laminate working electrode | SWV | 0.030 $\mu\text{M}$ | (47) |
| Paper-based sensor (carbon electrode) | SWV | 0.095 $\mu\text{M}$ | (48) |
| Boron-doped diamond (BDD) thin-film electrode | DPV | 0.05 $\mu\text{M}$ | (49) |
| -T-Macro | SWV | 0.51 $\mu\text{M}$ | (50) |
| -1.54T-CUA | | 1.0 $\mu\text{M}$ | |
| -CS/GNP 1.54T-CUA | | 1.6 $\mu\text{M}$ | |
| PANI/Au NPs/ITO | CV | 0.5 $\mu\text{M}$ | **The present work** |

**Figure 5.**
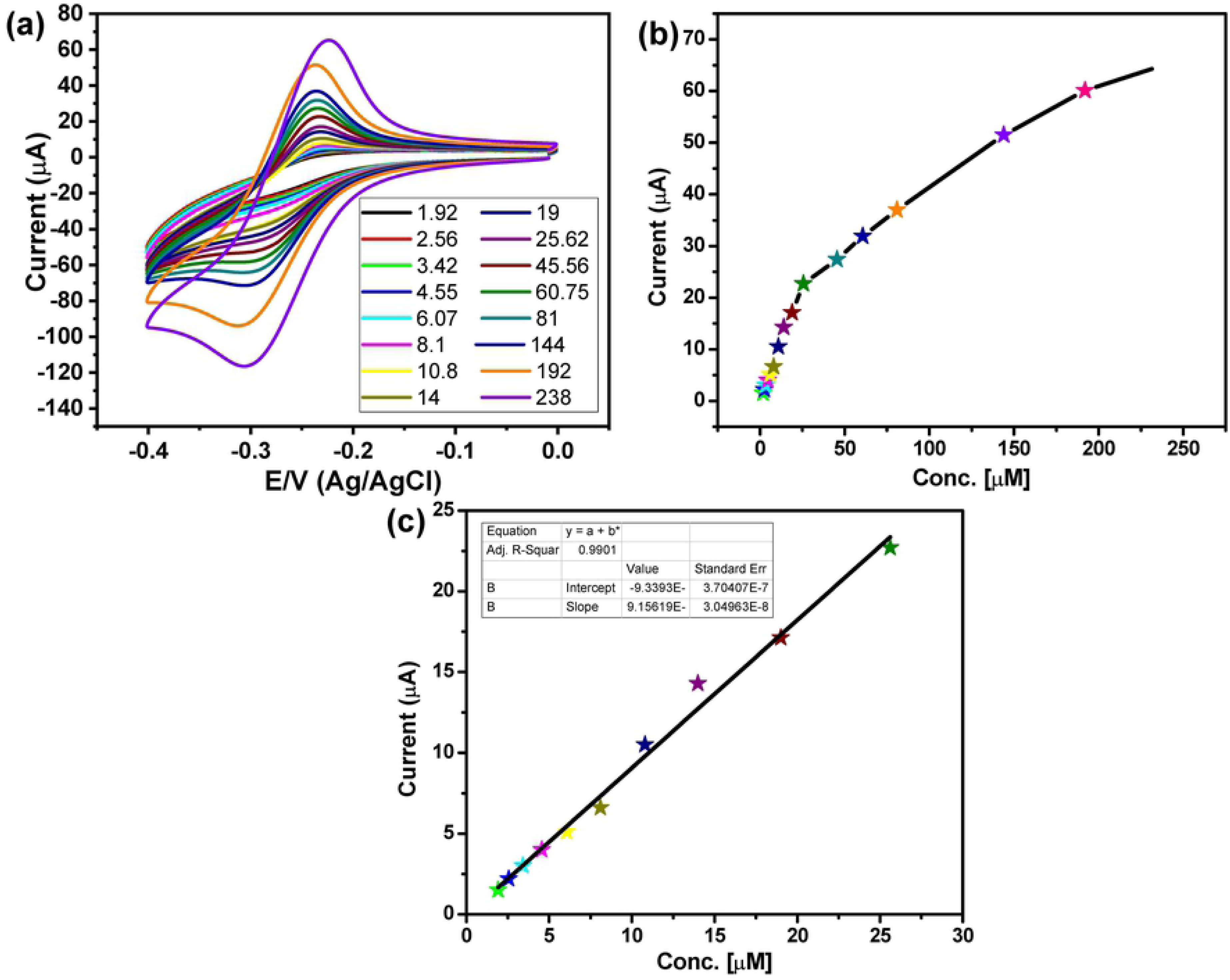
(a) Cyclic voltammograms of different concentrations of pyocyanin from 238 *µ*M to 1.9 *µ*M at scan rate 50 mV/sec, (b) relationship between the anodic current peaks and the pyocyanin concentration, and (c) linear relation between current peak and pyocyanin concentrations from 25.62 *µ*M to 1.9 *µ*M.

### 3.4. Selectivity of the developed sensor towards pyocyanin in the presence of different interferences

One of the concerns raised about the utility of the PANI/Au NPs modified ITO electrode for investigating the presence of *P. aeruginosa* in human samples is that there may be other molecules, which may interfere with electrode performance. It is always of great importance to achieve the highest selectivity and sensitivity towards pyocyanin in the presence of different interferences in clinical samples. Our study is focused on the selective detection of pyocyanin in presence of vitamin c, glucose and urea, which may present in clinical samples. **Figure 6** represented the CV of pyocyanin and interferences; there are no peaks of interferences in the potential window of pyocyanin. According to the obtained results, the selective detection of pyocyanin is applicable with high sensitivity. The results reported (**Figure 6**) showed a clear electrochemical fingerprint of pyocyanin, which was clearly observed when pyocyanin is measured among other redox-active compounds such as vitamin C, urea and glucose.

**Figure 6.**
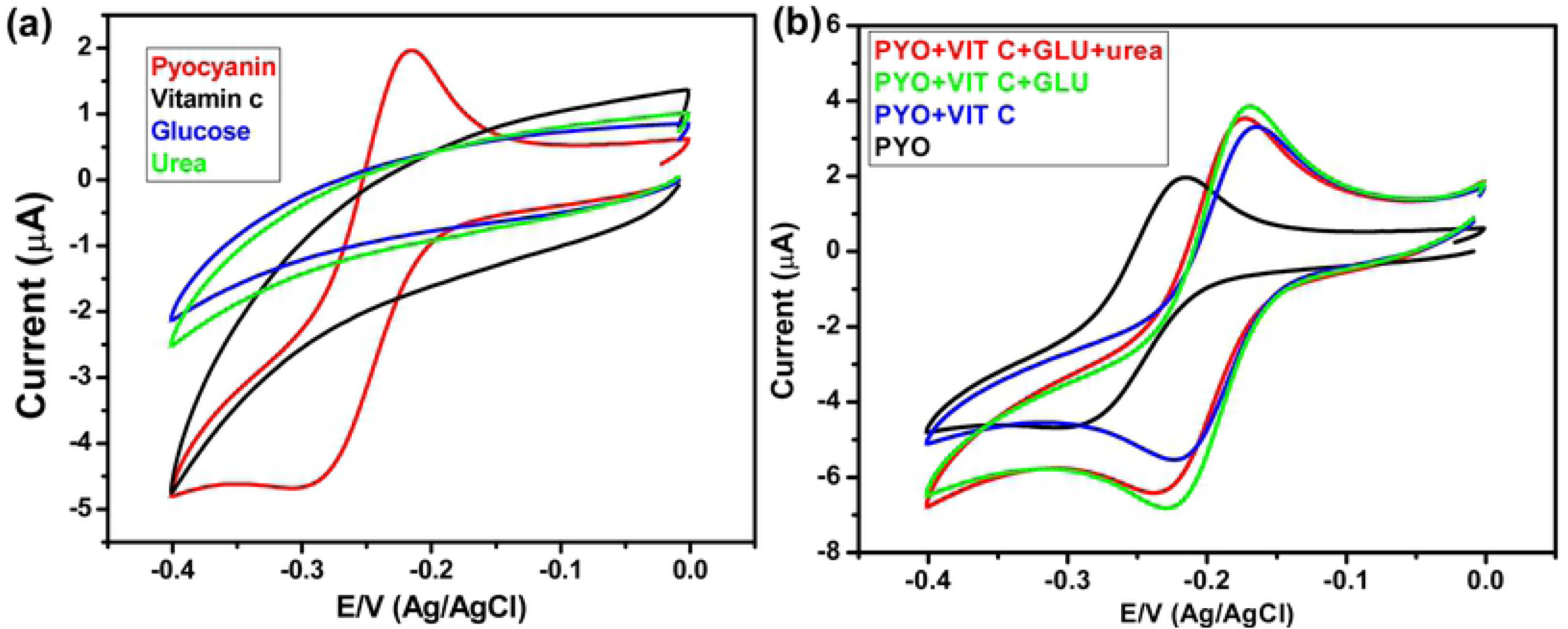
(a) CV of pyocyanin, vitamin c, glucose and urea, and (b) CV of pyocyanin and a mixture of pyocyanin, vitamin c, glucose and urea. interference of vitamin c, urea and glucose with the electrochemical detection of pyocyanin

### 3.5. Electrochemical detection of pyocyanin in Pseudomonas aeruginosa culture

In this study, samples from the *Pseudomonas aeruginosa* cultures were collected during log and stationary phase. The pyocyanin was released from *P. aeruginosa* culture and detected electrochemically by PANI/Au NPs modified ITO after 2,10 and 24 hours. The results in (**Figure 7**) show that no pyocyanin was initially produced in the culture after 2 hours of incubation as the OD at 600 nm was 0.1 so the culture was in the log phase. Pyocyanin could be detected after 10 hours as the culture reached the stationary phase. This result is consistent with the results obtained by Cabeen 2014 who reported that pyocyanin release is controlled by the quorum sensing system, which is not present in the early stage of a growing culture [**51**].

**Figure 7.**
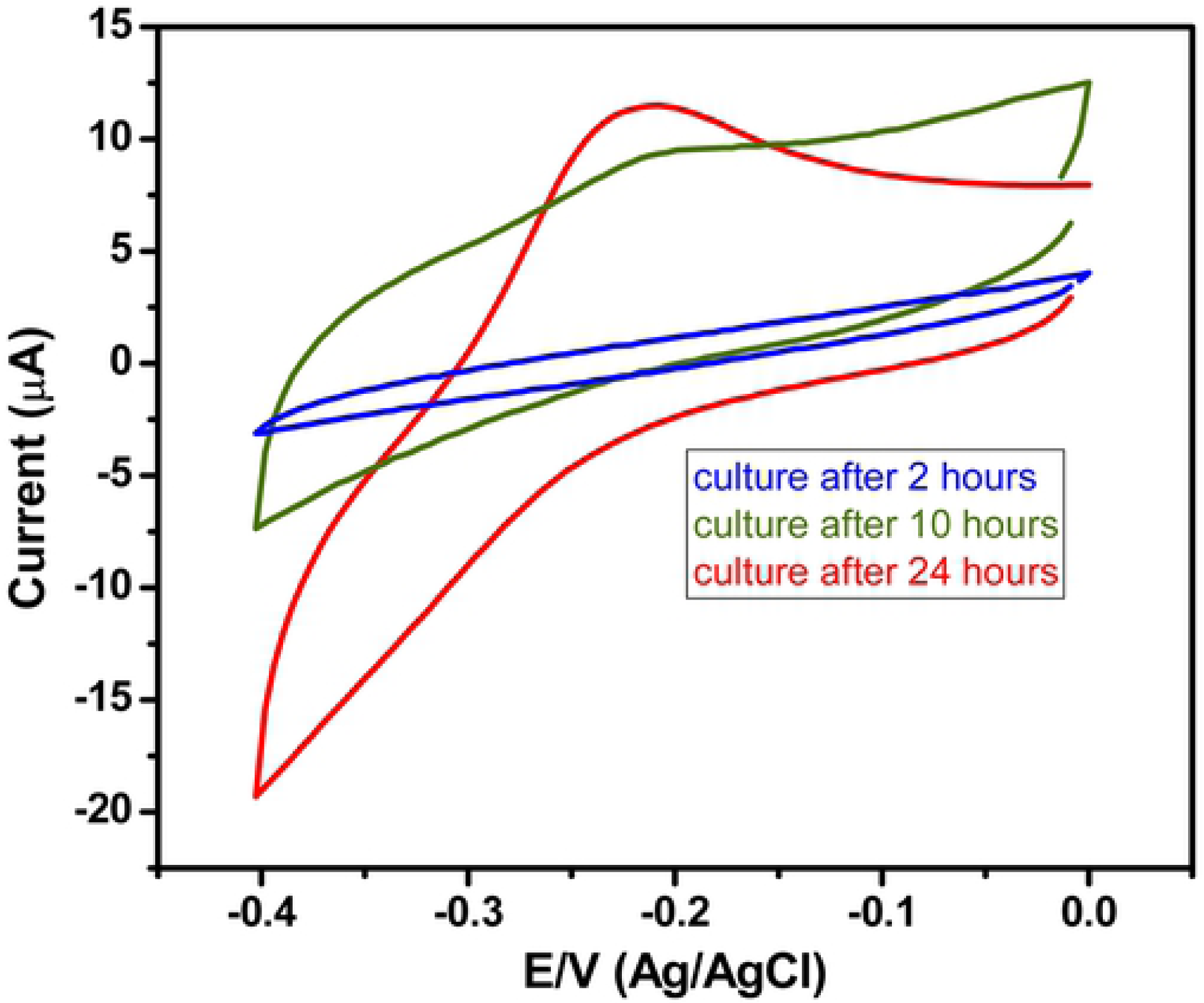
Electrochemical detection of pyocyanin in *P.aeruginosa* culture at 37 °C after 2, 10 and 24 hours of incubation

### 3.6. Electrochemical testing of bacterial cultures

The concern in this application was whether other bacterial pathogens produce redox-active molecules would interfere with the sensor’s response in the potential window of pyocyanin. To address this concern, a range of clinically-relevant bacterial pathogens (listed in **Figure 8**) were electrochemically measured after 24 hours of growth using the PANI/Au NPs modified ITO. The results in (**Figure 8**) indicated that the sensor demonstrated high selectivity towards pyocyanin. A notable oxidation peak at −0.23 V was originated from *P. aeruginosa* strain and there were no other bacteria that produce a redox-active peak in this potential window. The lack of a detectable peak from the other pathogens in the potential window further confirms that *P.aeruginosa* is the only bacterium producing redox-active molecules among the species tested and that the possibility of a false positive identification of *P.aeruginosa* using this method is unlikely.

**Figure 8.**
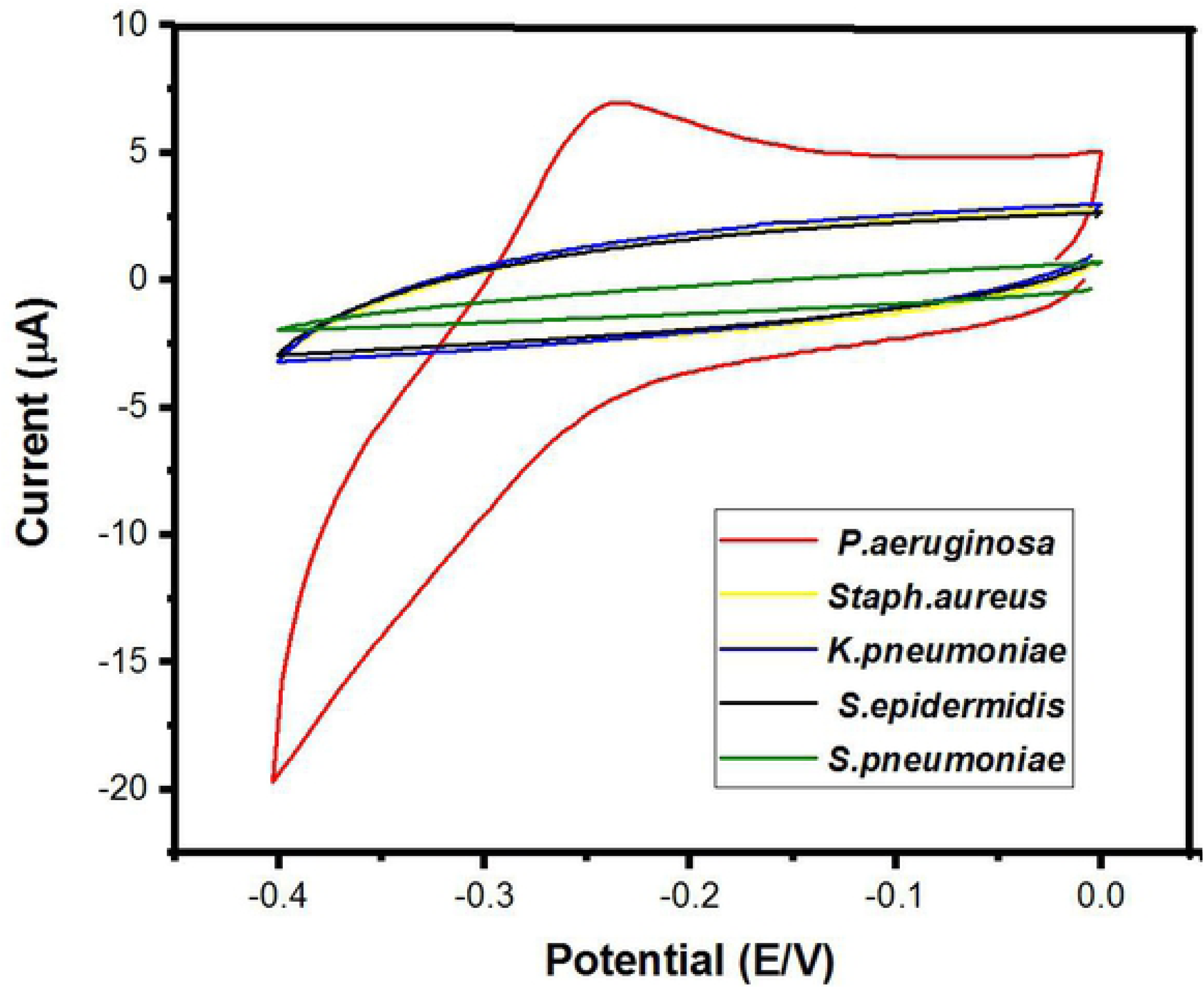
Cyclic voltammetry of different bacterial cultures after one day of growth at 37 °C.

### 3.7. Electrochemical measurements of clinical P. aeruginosa strains

Liquid cultures of clinical *P. aeruginosa* isolates were incubated at 37 °C for 24 hours with electrochemical measurements taken after incubation. All *P. aeruginosa* isolates were having a positive test result of electrochemical detection by PANI/Au NPs modified ITO electrode. The observed electrochemical peak (shown in **Figure 9**), due to pyocyanin oxidation at −0.20V, indicates the presence of *P.aeruginosa* in the sample. The negative control is LB broth growth media, which lacks the redox-active oxidation peak.

**Figure 9.**
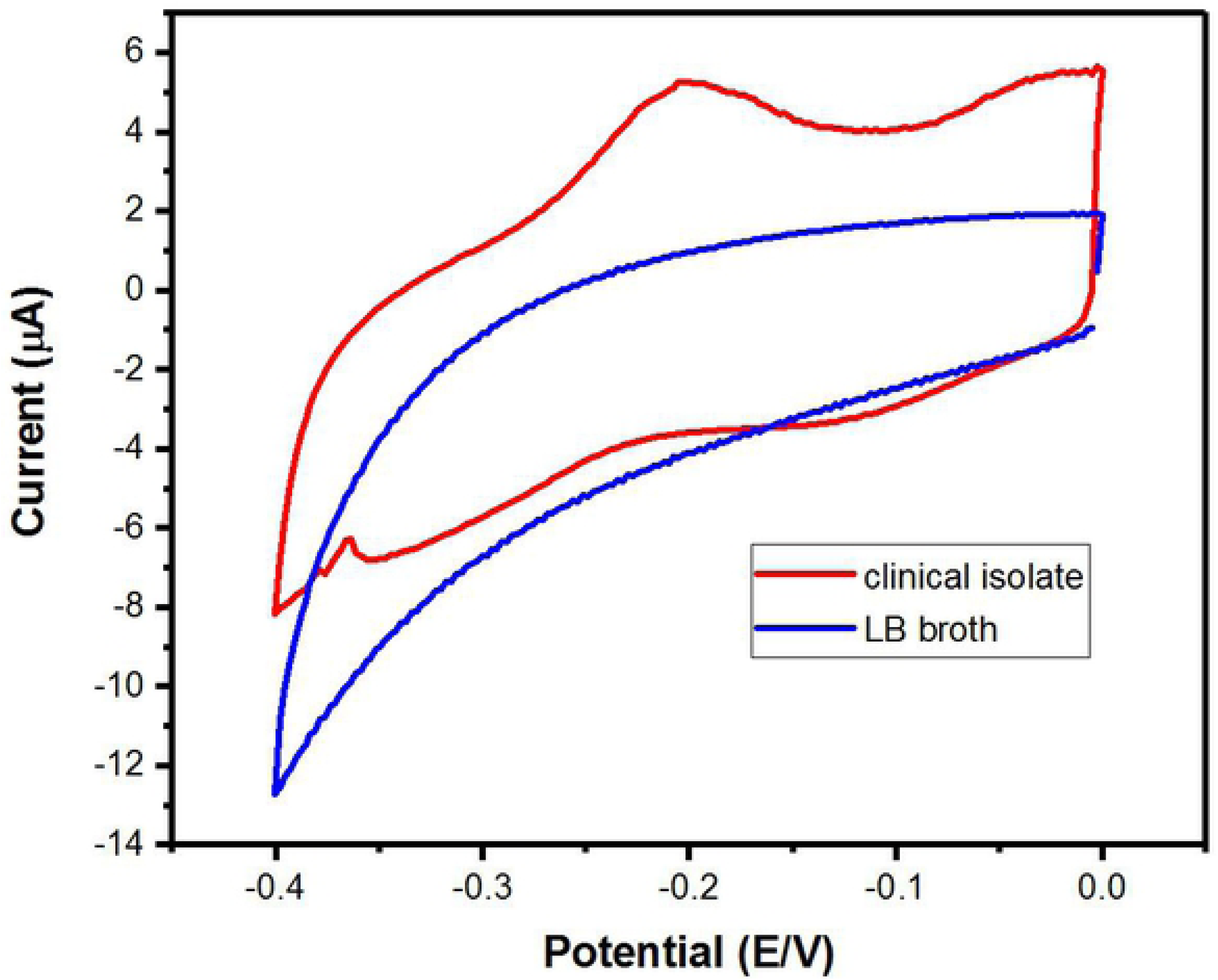
Cyclic voltammetry of a clinical isolate of *P. aeruginosa.*

## 4. Conclusions

In this work, we have fabricated a PANI/Au NPs modified ITO electrode based on the electrodeposition of Au NPs onto the ITO surface by using CV technique, followed by covering the surface with a layer of PANI. The prepared electrode showed 100% sensitivity, selectivity and a low detection limit for pyocyanin. The capability of the PANI/Au NPs modified ITO sensor to detect pyocyanin released in *P. aeruginosa* culture will aid in the fast precise detection of pyocyanin biomarker and diagnosis of *P. aeruginosa* infections specially in critically ill patients. Cosequently, this will achieve a rapid appropriate treatment and reduce the emergence of resistance made by empirical treatment.

